# The antioxidants esculetin and quercetin modulate differently redox balance in human NB4 leukemia cells

**DOI:** 10.1101/2024.02.28.578584

**Authors:** Virginia Rubio, Ana Isabel García-Pérez, Angel Herráez, Lilian Puebla, José C. Diez

## Abstract

The aim is to study the involvement of reactive oxygen species on redox homeostasis of acute promyelocytic leukemia cells using two antioxidants as models for tentative leukemia therapies. The NB4 leukemia cell line has increased reactive oxygen species (ROS) levels. Esculetin and quercetin are antioxidants that alter redox equilibrium, diminishing cell viability and inducing cell death in some cancer cells. The analysis of the role of reactive oxygen species on redox balance and apoptosis in NB4 leukemia cells was addressed by modulating redox balance with oxidants hydrogen peroxide (H_2_O_2_) and *tert*-butylhydroperoxide (*t*-BHP) and antioxidant compounds (esculetin and quercetin). Cell viability of NB4 cells was measured by flow cytometry using propidium iodide (PI). Colorimetric MTT (3-(4, 5-dimethylthiazol-2-yl)-2,5-diphenyltetrazolium bromide) gave an index of cell metabolism. Apoptosis was measured using Annexin V-FITC (fluorescein isothiocyanate coupled to annexin V). Intracellular ROS levels were detected using the fluorescent probes H_2_DCFDA (2’, 7’-dichloro-dihydrofluorescein diacetate) and DHE (dihydroethidine). Pre-treatment with quercetin prevented early loss of cell viability induced by either H_2_O_2_ or *t*-BHP. It reduced apoptosis produced by H_2_O_2_ and prevented that induced by *t*-BHP. Pre-treatment with esculetin, in contrast, increased apoptosis induced by H_2_O_2_ but reduced that produced by *t*-BHP. The superoxide anion content increased after incubation with either H_2_O_2_ or *t*-BHP, mainly in esculetin pretreatment. Esculetin, but not quercetin, prevented peroxide production by either H_2_O_2_ or *t*-BHP. Therefore, esculetin and quercetin regulate differentially redox balance in human NB4 human leukemia cells, presumably via different redox mechanisms, which opens up potential applications in antileukemia therapy.

## Introduction

Coumarins, commonly used as anticoagulants, are natural α-benzopyrones with an important therapeutic potential as anticancer, antioxidant and anti-inflammatory agents [Peng *et al*., 2013; Venugopala *et al*., 2013; Pereira *et al*., 2018]. They act as antioxidants by both diminishing ROS production and scavenging them.

Esculetin (6,7-dihydroxycoumarin) is a derivative of coumarin present in chicory as well as in many toxic and medicinal plants. Esculetin shows pro-apoptotic activity in leukemia cells and other tumoral cells by modulating redox balance [Kim *et al*., 2008]. In leukemia cell lines, esculetin inhibits cell cycle progression and shows a proapoptotic effect by activating caspase-3 and caspase-9, reducing Bcl2/Bax ratio without affecting p53 levels in NB4 cells [Rubio *et al*., 2014]. Esculetin also induces G_0_/G_1_ cell cycle arrest by suppressing cyclins D1 and D3 through a mechanism that blocks the Raf/MEK/ERK signalling pathway in HL-60 leukemia cell lines [Wang *et al*., 2019]. However, proapoptotic action of esculetin involves increased levels of phosphorylated ERK and phosphorylated Akt in NB4 cells [Rubio *et al*., 2014].

Quercetin (3,5,7,3’,4’ pentahydroxyflavone) has been studied as a potential therapeutic agent due to its effects on the suppression of many tumor-related processes. Quercetin induces cell death in leukemia cells by targeting epigenetic regulators affecting pro-apoptotic genes expression [Alvarez *et al*., 2018]. Quercetin reduced tumor growth in HL-60 xenografts accompanied by decreased expression of anti-apoptotic proteins, BCL-2, BCL-XL and MCL-1, and increased expression of the pro-apoptotic protein BAX with caspase-3 activation. Quercetin also mediated G_1_ phase cell cycle arrest in HL-60 xenografts [Calgarotto *et al*., 2018]. In K562 leukemia cells, quercetin promotes cell cycle arrest and induces apoptosis by modulation of the translational machinery and antioxidant defence systems [Alvarez *et al*., 2018; Liu *et al*., 2018]. Quercetin and other flavonoids directly interact with mitochondria causing respiratory chain inhibition that induces apoptosis by ROS generation and impairment of mitochondrial membrane potential [Altundag *et al*., 2018; Kim & Park, 2018; Cheng *et al*., 2019; Hassanzadeh *et al*., 2019; Brisdelli *et al*., 2020]. It may also affect the redox state by altering the intracellular glutathione content [Ramos & Aller, 2008; de Blas *et al*., 2016]. Quercetin also protects by the maintenance of cytosolic antioxidant enzymes activity (superoxide dismutase and catalase), protein sulfhydryl oxidation, and H_2_O_2_ production [de Lacerda *et al*., 2021].

The balance between the high production of reactive oxygen species (ROS) and decreased antioxidant defence systems leads to oxidative stress in cells [Dunning *et al*., 2013; Juránek *et al*., 2013; Huang *et al*., 2023]. Superoxide anion (O_2_^−^), hydrogen peroxide (H_2_O_2_) and the hydroxyl radical (OH^−^), the main intracellular ROS, are well known to regulate cell survival or death pathways. Increased ROS levels induce antioxidant enzymes in order to regulate intracellular redox balance [Dunning *et al*., 2013]. Therefore, alterations in this balance by both oxidants and antioxidants affect cell fate [Fiedor & Burda, 2014]. While ROS behave as oncogenic by triggering cell growth, changes in their intracellular levels can kill cancer cells [King & Yi, 2008; Hole *et al*., 2013].

Since tumor cells show high ROS levels and alterations in antioxidant systems, they are sensitive to changes in redox balance induced by oxidant or antioxidant compounds, which makes these feasible for cytotoxic applications as antitumor compounds [Juránek *et al*. 2013; Pisoschi & Pop, 2015; Zhu *et al*., 2015; Papiez *et al*., 2016].

NB4, an APL (acute promyelocytic leukemia) cell line, has been reported to have higher ROS levels than healthy cells [Lanotte *et al*., 1991; Hole *et al*., 2013]. These higher ROS levels promote cell proliferation by protecting cells from apoptosis [Hole *et al*., 2013]. Therefore, antioxidant compounds with ROS scavenger activity would induce apoptosis [King & Yi, 2008; Rubio *et al*., 2017a,b].

Esculetin, at low doses (1-25 μM), has been demonstrated to suppress H_2_O_2_-induced cell damage in human hepatoma HepG2 cells [Subramaniam & Ellis, 2011] and to protect against *t*-BHP oxidation in rat liver [Lin *et al*., 2000]. When administered in the diet (0.5% w/w) it protects against N-nitrosodiethylamine-induced hepatotoxicity in rats, and from DNA damage induced by 1,2-dimethylhydrazine in rat colon [Kaneko *et al*., 2007; Subramaniam & Ellis, 2011]. At higher doses (100-1000 μM), however, esculetin induces apoptosis in other cell lines such as adipocyte 3T3-L1 cells [Yang *et al*., 2006], human leukemia U937 cells [Park *et al*., 2008; Park *et al*., 2010] and human promyelocytic leukemia HL-60 cells [Wang *et al*., 2019]. A ROS-mediated mitochondrial-dependent apoptotic pathway may be relevant in these processes [Kim *et al*., 2008; King & Yi, 2008]. Our group found that esculetin produces significant and dose-dependent apoptosis in NB4 cells. considering a role of ROS in this cell death mechanism [Rubio *et al*., 2014]. We also showed that esculetin acts synergistically with H_2_O_2_ to decrease cell viability and metabolic activity, as well as to increase apoptosis at early times when both are added simultaneously [Rubio *et al*., 2017a]. In contrast, esculetin neutralised the effects of the oxidant *t*-BHP when both were supplied simultaneously and prevented apoptosis of human leukemia NB4 cells. Therefore, it prevents the oxidant effect of *t*-BHP but not the effect of H_2_O_2_ on NB4 cells [Rubio *et al*., 2017a]. On the other hand, quercetin and many polyphenols selectively induce apoptosis in leukemia cells and potentiate the efficacy of conventional anti-cancer drugs by reducing intracellular ROS [Fantini *et al*., 2006; Jeon *et al*., 2019].

To understand the apoptotic mechanism, related to ROS, of esculetin and quercetin, in this study we have analyzed the role of esculetin, in comparison with quercetin, on apoptosis induced by the exogenous ROS generators *tert*-butylhydroperoxide (*t*-BHP) or H_2_O_2_. *t*-BHP is a short chain analogue of lipid hydroperoxides, similar to peroxidised fatty acids, which has been widely used to induce oxidative stress in a variety of cells by increasing ROS and decreasing GSH [Lanotte *et al*., 1991; Kaneko *et al*., 2007]. H_2_O_2_ is generated under oxidative stress conditions modulating intracellular processes [Rincheval *et al*., 2013]. To study intracellular ROS levels at short time period after oxidant administration, NB4 cells were pre-treated with either esculetin or quercetin and subsequently exposed to either *t*-BHP or H_2_O_2_.

## Materials and methods

### Reagents

Esculetin (6,7-dihydroxycoumarin, 98% purity) and quercetin were obtained from Sigma-Aldrich (Steinheim, Germany) and prepared as 196 mM and 100 mM stock solutions respectively in dimethyl sulfoxide (DMSO) and stored at −20°C. H_2_O_2_ (hydrogen peroxide) and *t*-BHP (*tert*-butyl hydroperoxide) were purchased from Sigma-Aldrich (Steinheim, Germany) and prepared as 0.88 M and 0.078 M stock solutions respectively in distilled water at the time of use. Fluorescent probes dihydroethidine and 2’,7’-dichlorodihydrofluorescein diacetate were obtained from Molecular Probes (Eugene, Oregon, USA).

### Cell culture

The human NB4 leukemia cell line was obtained by our group from the American Type Culture Collection [Galeano *et al*., 2005; Sancho *et al*., 2007]. Cell line was maintained in culture at a density of 3×10^5^ cells/ml in RPMI medium (Gibco-Life Technologies) supplemented with 10% fetal bovine serum (FBS), 1% penicillin+streptomycin and 0.02 mg/ml gentamicin at 37°C in a humidified 5% CO_2_ atmosphere [Galeano *et al*., 2005; Sancho *et al*., 2007].

### Cell treatments

100 μM esculetin or 25 μM quercetin were added to NB4 cells (5×10^5^ cells/ml) for 30 min or 2h. Afterwards, the cells were incubated for 1 h with the oxidants, 1 mM H_2_O_2_ or 250 μM *t*-BHP. Appropriate controls were run in parallel.

### Cell viability study

NB4 cell viability was determined by measuring the permeability to propidium iodide (PI) by flow cytometry. After treatments, 2.5×10^5^ cells were washed with 500 μl phosphate-buffered saline (PBS) and resuspended in 300 μl PBS. Then, 15 μl propidium iodide (Calbiochem) were added and the fluorescence was measured using a Becton Dickinson FACScalibur flow cytometer (San José, CA, USA) [Rubio *et al*., 2014].

### Cell metabolic viability

Cell metabolic viability was measured by the colorimetric MTT (3-(4,5-dimethylthiazol-2-yl)-2,5-diphenyltetrazolium bromide) assay kit (Roche). After treatments, the cells seeded in 96-well microplates were incubated with 10 μl MTT Labelling Reagent for 4 h and then 100 μl of the solubilisation solution were added. A microplate reader rendered measurements of absorbance which correlate with the number of viable cells.

### Analysis of apoptosis by Annexin-V-FITC cytometry

Apoptosis was measured by the presence of phosphatidylserine on the outer side of the cell membrane, using Annexin V-FITC (fluorescein isothiocyanate coupled to annexin V) Apoptosis Detection Kit (BioVision). After treatments, 2.5×10^5^ cells were centrifuged at 1200 rpm for 5 min and incubated with 500 µl 1× Annexin V Binding Buffer and 1 µl Annexin V-FITC for 5 min at room temperature in the dark. Then, 10 μl of PI were added and the apoptotic cells measured by fluorescence using a Becton Dickinson FACScalibur flow cytometer. WinMDI 2.8 software was used for analysis of results [Rubio *et al*., 2014].

### Measurement of intracellular ROS levels

Intracellular ROS levels were detected using the fluorescent probe H_2_DCFDA (2’, 7’-dichlorodihydro-fluorescein diacetate), a non-fluorescent molecule which accumulates intracellularly and reacts with reactive oxygen species, especially hydrogen peroxide, becoming green fluorescent 2’,7’-dichlorofluorescein (DCF). After treatment, incubation of the cells was carried out with 10 μM H_2_DCFDA for 30 minutes at 37ºC, and the cells were washed with PBS. Finally, fluorescence intensity was measured using the FACScalibur flow cytometer.

To measure intracellular superoxide, cells were incubated with 2 μM DHE (dihydroethidine) during the final 15 min of the last treatment. Fluorescence intensity was measured by flow cytometry [Rubio *et al*., 2014].

### Statistical analysis

Data are displayed as the mean ± standard error of at least three independent experiments. The differences between the control and treated groups were determined using the ANOVA test and a p<0.05 value was considered statistically significant. In the figures, the asterisk * indicates comparison between treated and control untreated samples. The symbol + (effect of the oxidant) is used for comparison between samples treated first with esculetin or quercetin and then with H_2_O_2_ or *t*-BHP, versus samples treated exclusively with esculetin or quercetin. The symbol # (effect of the antioxidant) is used for comparison between samples treated first with esculetin or quercetin and then with H_2_O_2_ or *t*-BHP, versus samples treated exclusively with H_2_O_2_ or *t*-BHP. In all cases, one, two or three symbols indicate p <0.05, p <0.01 and p <0.001, respectively.

## Results

### Cell viability and metabolic activity in the presence of H_2_O_2_ or tBHP of NB4 cells pre-treated with esculetin or quercetin

Previous studies in our group have shown that the coumarin esculetin and the flavonoid quercetin exert different antioxidant effects on NB4 cells, probably due to the mechanisms by which they regulate cellular redox balance [Rubio *et al*., 2018]. Here, we comparatively studied the counterbalance effects of quercetin or esculetin in preventing oxidation caused by H_2_O_2_ or *t*-BHP in human NB4 leukemia cells during the first hour of treatment. According to our previous studies, the lowest concentration of esculetin capable of decreasing NB4 cell viability was 100 μM when applied for as long as 19 h [Rubio *et al*., 2014]. On the other hand, quercetin also reduced NB4 cell viability at 25 μM for 24 h [Rubio *et al*., 2019]. Therefore, these concentrations were selected to study the influence of both antioxidants on ROS regulation. The H_2_O_2_ and *t*-BHP concentrations were the same as those used in our previous studies with esculetin [Rubio *et al*., 2017a,b].

Human leukemia NB4 cells were pre-incubated with either 100 μM esculetin or 25 μM quercetin for 30 min or 2 h. Subsequently, the cells were treated with 1 mM H_2_O_2_ or 250 μM *t*-BHP for 1 h. Cell viability was analyzed by measuring the permeability to propidium iodide using flow cytometry, whereas cell metabolic activity was ascertained by the MTT assay.

As can be observed in Figure 1, pretreatment with esculetin did not affect NB4 cell viability, whereas treatment with either H_2_O_2_ or *t*-BHP, alone, for 1 hour clearly reduced it. When esculetin preceded *t*-BHP, it abolished the decrease in cell viability. On the contrary, when cells were pre-treated with esculetin for different time-periods and subsequently treated with H_2_O_2,_ an early reduction in cell viability was observed, as compared to untreated control cells, indicating that esculetin did not protect against the effects of H_2_O_2,_ mainly at very short time-periods (Figure 1). In summary, treatments with esculetin prevent loss of cell viability induced by *t*-BHP but not by H_2_O_2_.

**Figure 1.**
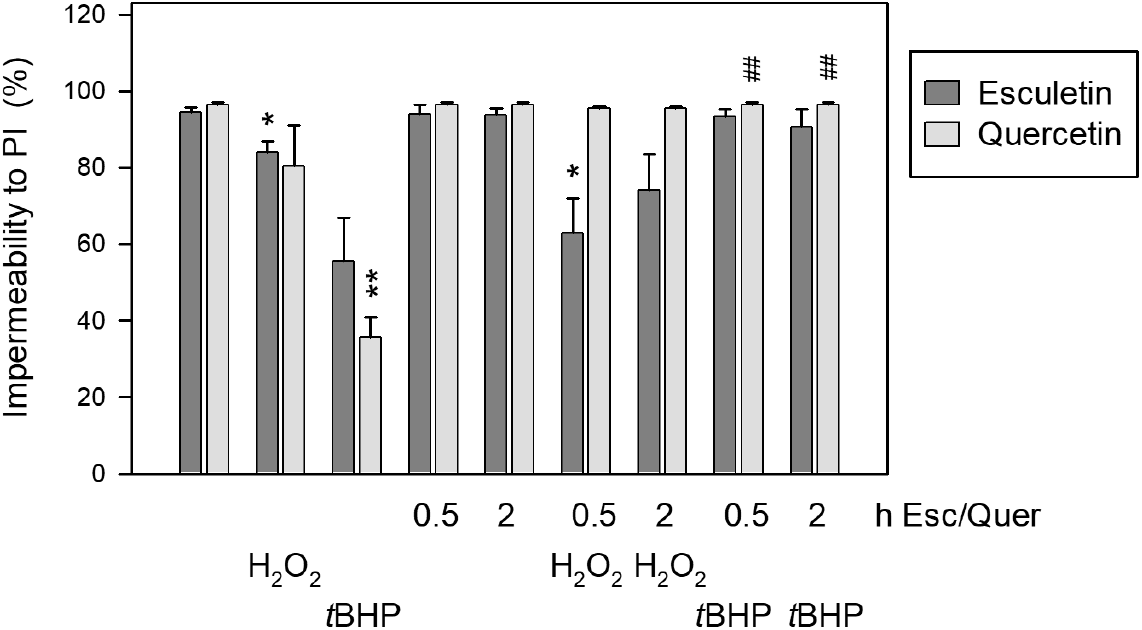
NB4 cell viability determined by flow cytometry as the impermeability to PI. 5×10^5^ cells/ml were pre-treated with either 100 μM esculetin or 25 μM quercetin for 30 min or 2 hours. Then, cells were supplemented with either 1 mM H_2_O_2_ or 250 μM *t*-BHP for 1 hour. The results represent the mean ± SEM of three independent experiments. The asterisk **\*** indicates statistical significance comparing treated to control untreated samples. Symbol **#** reflects the effect of the antioxidant (see the Methods section for more details). In all cases, one, two or three symbols indicate p<0.05, p <0.01 and p <0.001, respectively.

This was not the case when NB4 cells were pretreated with quercetin since subsequent treatments with H_2_O_2_ or *t*-BHP did not affect cell viability (Figure 1). Thus, pre-treatment with quercetin prevents loss of cell viability induced by either H_2_O_2_ or *t*-BHP.

Figure 2 shows the metabolic viability of NB4 cells treated under the same conditions as above. Esculetin or quercetin had a relatively low effect on NB4 metabolic activity, even after incubation for 2 hours. Independent treatments of NB4 cells with either H_2_O_2_ or *t*-BHP reduced the metabolic viability. Pre-treatment with esculetin, however, showed a partial protection against reduction of metabolic activity when caused by *t*-BHP but not by H_2_O_2_ (Figure 2). Quercetin pre-treatments slightly protected against the loss of metabolic activity caused by H_2_O_2_ albeit less than that caused by *t*-BHP (Figure 2).

**Figure 2.**
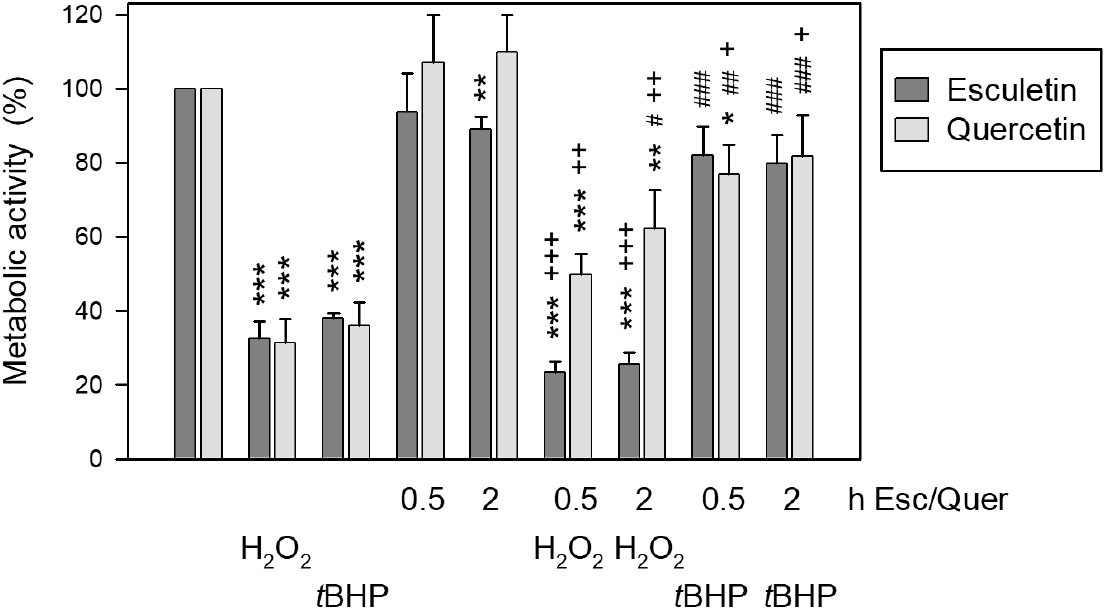
Metabolic viability of NB4 cells measured by the MTT assay. 5×10^5^ cells/ml were pre-treated with either 100 μM esculetin or 25 μM quercetin for 30 min or 2 hours. Then, cells were supplemented with either 1 mM H_2_O_2_ or 250 μM t-BHP for 1 hour. The results represent the mean ± SEM of three independent experiments. The asterisk **\*** indicates statistical significance comparing treated to control untreated samples. Symbol **+** reflects the effect of the oxidant. Symbol **#** reflects the effect of the antioxidant (see the Methods section for more details). In all cases, one, two or three symbols indicate p<0.05, p <0.01 and p <0.001, respectively.

Thus, pre-treatments with esculetin partly prevent reduction of metabolic activity induced by *t*-BHP but not by H_2_O_2_, while quercetin protects from H_2_O_2_ less than from *t*-BHP.

### Apoptosis and necrosis in the presence of H_2_O_2_ or tBHP of NB4 cells pre-treated with esculetin or quercetin

We next studied apoptosis and necrosis of NB4 cells treated with either 100 μM esculetin or 25 μM quercetin for two different time-periods (30 min or 2 h) and then incubated with either 1 mM H_2_O_2_ or 250 μM *t*-BHP for 1 h (Figure 3). Esculetin pretreatments seemed to prevent early apoptosis induced by *t*-BHP but not by H_2_O_2_. Quercetin treatments resulted in very low levels of apoptosis even after subsequent incubations with H_2_O_2_ or *t*-BHP. Necrosis of NB4 cells was not observed in any of the cases except slightly in that of *t*-BHP treatment.

**Figure 3.**
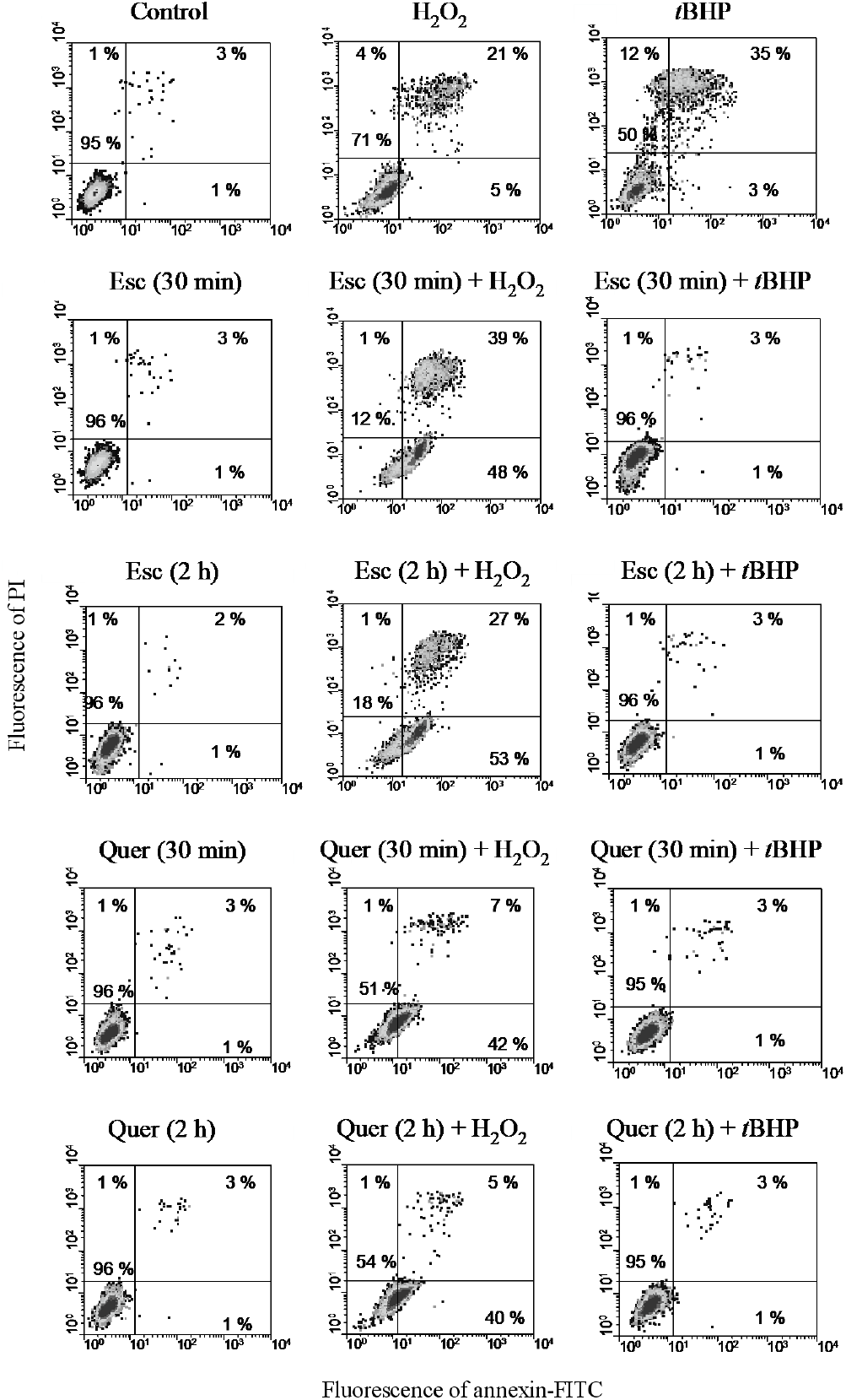
Apoptosis and necrosis of NB4 cells measured by flow cytometry after staining with FITC-conjugated annexin and PI. 5×10^5^ cells/ml were pre-treated with either 100 μM esculetin or 25 μM quercetin for 30 min or 2 hours. Then, cells were supplemented with either 1 mM H_2_O_2_ or 250 μM t-BHP for 1 hour. The results show one representative experiment out of three previous experiments.

### ROS production in the presence of H_2_O_2_ or tBHP in NB4 cells pre-treated with esculetin or quercetin

In order to consider a tentative relationship between the above results and ROS levels, we first used the specific probe DHE to detect superoxide anion by flow cytometry under the different conditions. Esculetin or quercetin, independently, did not significantly affect the superoxide levels in NB4 cells. Interestingly, pre-treatments with either esculetin or quercetin and subsequent incubation with 1 mM H_2_O_2_ increased superoxide levels (Figure 4). However, pre-treatment with either of the two antioxidants attenuated the superoxide induced by *t*-BHP.

**Figure 4.**
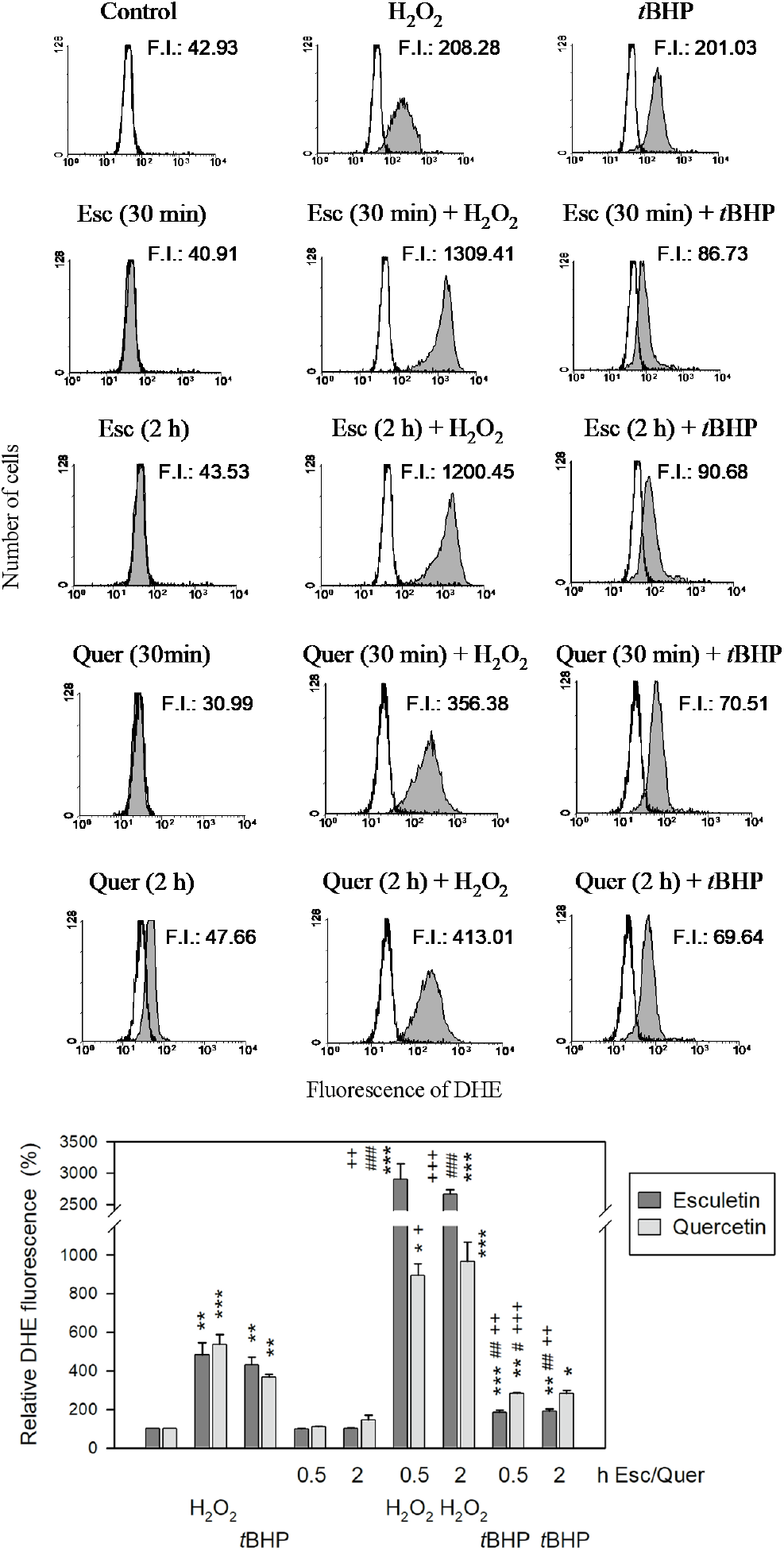
Intracellular superoxide levels in NB4 cells measured by flow cytometry with DHE, a specific probe. 5×10^5^ cells/ml were pre-treated with either 100 μM esculetin or 25 μM quercetin for 30 min or 2 hours. Then, cells were supplemented with either 1 mM H_2_O_2_ or 250 μM *t*-BHP for 1 hour. During the last 15 min, 2 μM DHE was added. The upper panel shows the results of one representative experiment out of three. The lower panel shows the mean ± SEM of three independent experiments. The asterisk **\*** indicates statistical significance comparing treated to control untreated samples. Symbol **+** reflects the effect of the oxidant. Symbol **#** reflects the effect of the antioxidant (see the Methods section for more details). In all cases, one, two or three symbols indicate p<0.05, p <0.01 and p <0.001, respectively.

We next sought to determine whether or not there was a relationship between cytotoxicity and peroxide levels using H2DCFDA as a fluorescent probe to detect peroxides by flow cytometry (Figure 5). Treatments with H_2_O_2_ or *t*-BHP produced increments in peroxide anion level in NB4 cells that were prevented by esculetin. On the contrary, quercetin pre-treatment increased the peroxide content produced by *t*-BHP alone.

**Figure 5.**
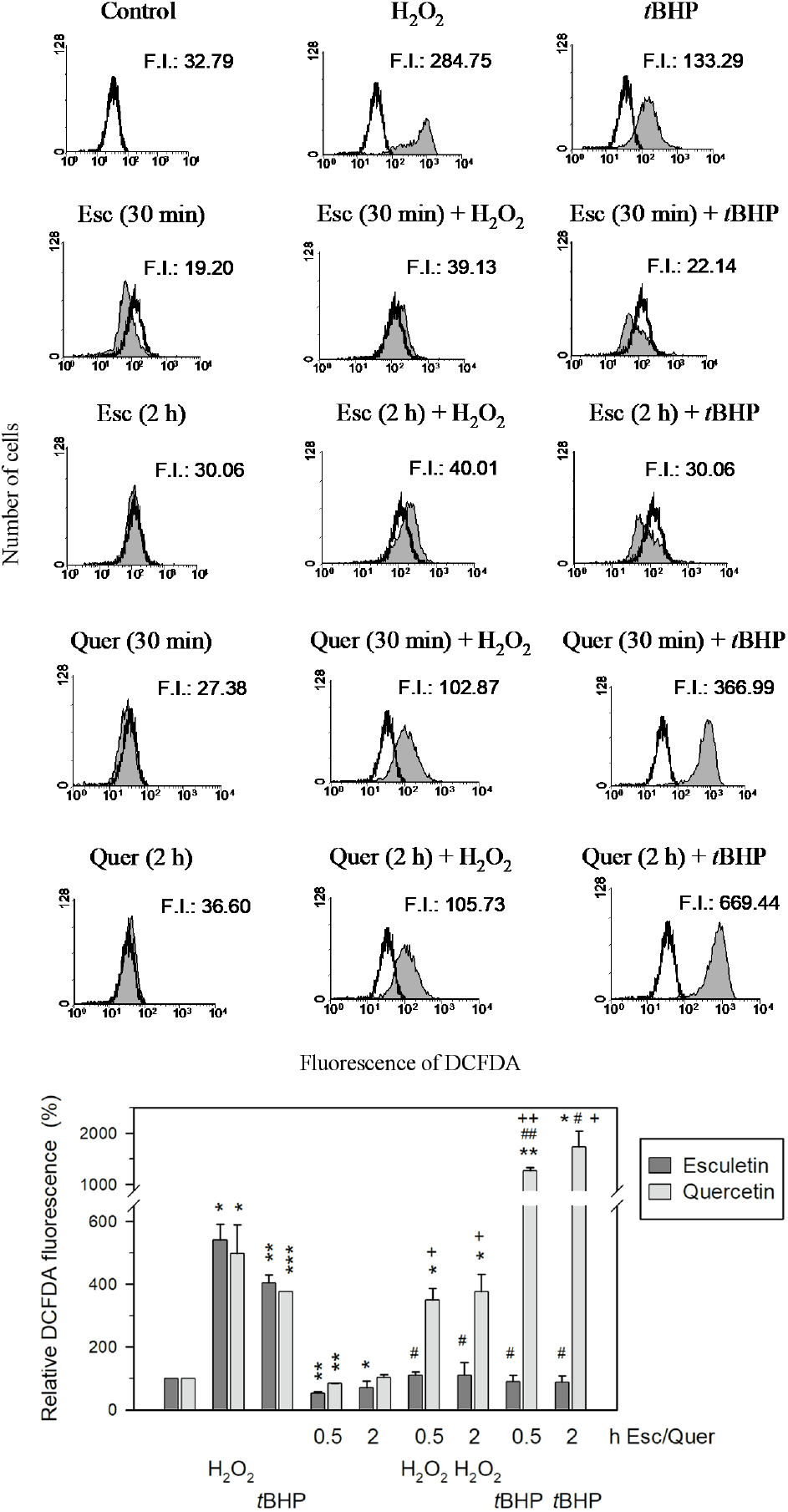
Intracellular peroxide levels in NB4 cells, measured by flow cytometry with the specific probe H_2_DCFDA. 5×10^5^ cells/ml were pre-treated with either 100 μM esculetin or 25 μM quercetin for 30 min or 2 hours. Then, cells were supplemented with either 1 mM H_2_O_2_ or 250 μM *t*-BHP for 1 hour. During the last 30 min, 2 μM H_2_DCFDA was added. The upper panel shows the results of one representative experiment out of three. The lower panel shows the mean ± SEM of three independent experiments. The asterisk **\*** indicates statistical significance comparing treated to control untreated samples. Symbol **+** reflects the effect of the oxidant. Symbol **#** reflects the effect of the antioxidant (see the Methods section for more details). In all cases, one, two or three symbols indicate p<0.05, p <0.01 and p <0.001, respectively.

Altogether, our results show different early effects of the antioxidants esculetin and quercetin on NB4 leukemia cells, indicating different mechanisms of action of these two compounds; this is suggestive of differential applications of both compounds for antitumor therapy.

## Discussion

Our previous studies in human leukemia NB4 cells have demonstrated that esculetin is able to scavenge ROS when added in a cotreatment with the oxidants H_2_O_2_ and [Rubio *et al*., 2014]. It has been proposed that ROS reduction is followed by cell growth arrest and apoptosis, via an intrinsic mitochondrial pathway [Muller *et al*., 2007]. Esculetin acts early (15 min) and synergistically with H_2_O_2_, decreasing cell viability and increasing apoptosis in NB4 cells [Rubio *et al*., 2017a,b]. In contrast, esculetin neutralizes early the oxidative effects of *t*-BHP and hence protects against apoptosis [Rubio *et al*., 2017a].

To improve our knowledge on the different apoptotic mechanisms of esculetin in the presence of these oxidants, the present study was also carried out with the known antioxidant polyphenol quercetin for comparative purposes. Quercetin has shown a protective effect against apoptosis in PC-12 cells by reducing ROS levels, malondialdehyde and lipoperoxidation of the cell membrane [Bao *et al*., 2017]. It also protects IPEC-J2 enterocytes from oxidative damage and apoptosis induced by H_2_O_2_ [Chen *et al*., 2018].

To determine whether the antioxidant activity of esculetin or quercetin can modulate differently NB4 cell apoptosis based on changes in redox equilibrium, we increased the oxidative stress by addition of some oxidants.

We carried out short-term treatments with esculetin or quercetin before incubating the cells with the oxidants *t*-BHP or H_2_O_2_. Treatment of NB4 cells with *t*-BHP for 1 h reduced both cell viability and metabolic activity and induced apoptosis. Pre-treatments with esculetin or quercetin, however, protected these cells from an early tBHP-induced apoptosis. The addition of *t*-BHP to NB4 cells affects ROS levels by increasing superoxide anion content and peroxides (Figures 4 and 5), but this is early offset by prior addition of esculetin. Pre-treatment with quercetin early decreases superoxide anion content (Figure 4) whereas it increases the levels of peroxides induced by *t*-BHP in NB4 cells (Figure 5). These results indicate a different early mechanism of ROS regulation by esculetin or quercetin with respect to the protection against peroxides.

In the presence of H_2_O_2_ (1 mM), both cell viability and metabolic activity were reduced and cells underwent apoptosis. Pre-treatment with esculetin did not protect against these early H_2_O_2_-induced effects, and actually increased the apoptosis produced by H_2_O_2_ (Figures 1-3). In contrast, pretreatment with quercetin prevented the early loss of cell viability although it did not fully restore metabolic activity to control levels and reduced apoptosis induced by H_2_O_2_. Hence, these antioxidants exert different early protection against H_2_O_2_ in NB4 cells.

Pre-treatment with either esculetin or quercetin does not protect against the early increased H_2_O_2_-induced superoxide, indicating a similar mechanism of action for both antioxidants. Esculetin, alone, significantly reduced the early increase of peroxides induced by H_2_O_2_ alone to levels close to those of control cells. By contrast, pre-treatment with quercetin did not lead to a significant reduction of peroxide early induced by H_2_O_2_.

One could conclude that both antioxidants do not behave as superoxide scavengers. There is a difference, however, regarding peroxides: while esculetin scavenged peroxides induced by H_2_O_2_ or *t*-BHP, quercetin did not exert any protection.

Thus, esculetin seems to be adequate in preventing exogenous oxidation via activation of the antioxidant response.

Some studies on the activity of esculetin have shown bioavailability of the antioxidant [Lin *et al*., 2000]. This gave rise to inhibitory activity of oxidative damage in liver [Lin *et al*., 2000]. Additionally, Wang *et al*. [2015] demonstrated antiproliferative activity on hepatocellular carcinoma in vivo, depending on mitochondrial dependent-apoptosis.

Similarly, in vivo antitumor action of quercetin has been also demonstrated in several animal models including hepatocarcinoma [Hashemzaei *et al*., 2017; Fernández-Palanca *et al*., 2019] showing bioavailability of the antioxidant.

Previous studies in our group with both antioxidants have shown that quercetin inhibits the antioxidant response mediated by Nrf2 whereas this effect is activated by esculetin [Rubio *et al*., 2018].

Changes in apoptosis factors (Bax, Bcl2) or in phosphorylation of kinases such as Erk can also be present in prevention of the observed ROS changes as an effect of these antioxidants, at least in the case of esculetin [Rubio *et al*., 2014].

Moreover, these effects on apoptosis factors may also be observed in the case of quercetin, as it has been shown in H_2_O_2_-induced apoptosis in intestinal porcine enterocytes [Chen *et al*., 2018].

The involvement of factors such as p53, Bax, Bcl2 and others, as well as some signalling kinases (Erk, etc.), on the differential modulation by these antioxidants of redox balance in the conditions here studied are now under current investigation.

These results support a distinct early mechanism of ROS regulation by the antioxidants esculetin or quercetin, which could be used to potentiate apoptosis of leukemia NB4 cells and must be taken into account for different therapy applications of modulation of redox balance in tumor cells.

## Conclusions

Esculetin increases H_2_O_2_-induced apoptosis but significantly reduces that produced by *t*-BHP. Quercetin reduces H_2_O_2_-induced apoptosis and wholly protects against apoptosis induced by *t*-BHP. Both esculetin and quercetin lead to a reduction of superoxide content in cells treated with *t*-BHP. Esculetin but not quercetin reduces peroxide production by H_2_O_2_ or *t*-BHP.

Our results show differential redox regulation in human NB4 leukemia cells by two different antioxidant compounds, esculetin or quercetin, presumably via different mechanisms, which opens up the possibility of potential applications of regulation of cell redox balance in cancer therapy.

## Acknowledgements

Financial support of this work was in part by research grants FISS PI060119, CCG10-UAH/SAL-5966 and UAH2011/BIO-006. We also want to thank Isabel Trabado for her technical assistance in cytometry analyses (C.A.I. Medicina-Biología. Unidad de Cultivos. Universidad de Alcalá).

